# A Conservative Approach for Describing Cancer Progression

**DOI:** 10.1101/2022.06.11.495730

**Authors:** Nicolò Rossi, Nicola Gigante, Nicola Vitacolonna, Carla Piazza

## Abstract

The field of tumor phylogenetics focuses on studying the differences within cancer cell populations and many efforts are done within the scientific community to build cancer progression models trying to understand the heterogeneity of such diseases. These models are highly dependent on the kind of data used for their construction and, as the experimental technologies evolve, it is of major importance to exploit their peculiarities. In this work we describe a cancer progression model based on Single Cell DNA Sequencing data. When constructing the model, we focus on tailoring the formalism on the specificity of the data, by defining a minimal set of assumptions to reconstruct a flexible DAG structured model, capable of identifying progression beyond the limitation of the infinite site assumption. We provide simulations and analytical results to show the features of our model, test it on real data, show how it can be integrated with other approaches to cope with input noise. Moreover, our framework can be exploited to produce simulated data that follows our theoretical assumptions. Finally, we provide an open source R implementation of our approach that is publicly available on BioConductor.

## 1 Introduction

Cancer, one of the primary causes of death in developed countries, is a genetic disease where mutations change the behavior of some body cells inducing an out of control proliferation, with effects on the host comparable to those of a parasitic entity. However, tumors are a complex class of diseases varying both at the macroscopic level (e.g., tumor location and size) and at the microscopic level (e.g., genetic asset and gene expression). Current models represent such genetic drift as an evolutionary process [1], albeit with its own peculiarities. According to such a view, a tumor originates from a single cell and progresses by acquiring genetic variability, and therefore giving rise to several genetically distinct and relatively unstable cell populations called *clones*, competing (or possibly cooperating) for the limited resources in their micro-environment. Several tumor evolution models have been proposed to explain such intratumor heterogeneity [2], and they aim to become powerful tools for understanding cancer progression and helping design effective treatments.

Exploiting this evolutionary perspective, several *tumor phylogenetic* techniques and methods have been developed over the years, either by adapting computational approaches used in biology to reconstruct species evolution or by creating newer models, specifically crafted for this context [3], [4]. Independently from the chosen strategy, it is possible to define three main categories of cancer data [3]:

- *cross-sectional* methods, that combine samples from different tumors of different patients;
- *regional bulk* methods, where samples from different tumor sites of a single patient are collected; and
- more recently, *single-cell* methods, which analyze genomic data sequenced from single cells originating from a single tumor site [5].

Cross sectional methods have shown how tumors tend to be quite diverse when considering primary site classification (see, e.g., [3] for an overview). To investigate this heterogeneity, *Single-Cell Sequencing* (*SCS*) samples are potentially the most useful because they allow the direct observation of instances of clones subpopulations, but they are also the hardest to collect, and the scarcest among the currently available datasets. Nevertheless, as genomic sequencing becomes cheaper and more accurate, SCS datasets are becoming easier to produce and collect, and suitable computational methods are needed for their exploitation.

In this paper, we describe a method to extract probabilistic models, which we call *Cancer Progression Markov Chains* (*CPMC*), from DNA SCS datasets, which describe the mutational history of the cells of a sampled tissue. CPMC are a particular kind of *Discrete Time Markov Chains* (*DTMC*) tailored to our use case, with useful mathematical properties. We first show that the kind of SCS datasets currently available can be described by infinite source models (called *generators*). Then, we introduce a realistic assumption on the process underlying the mutational events, and we show that, under such hypothesis, the solution to our inference problem is unique. An algorithmic method to find such a solution is described. When the uniqueness is not guaranteed we define an heuristic for inferring one of the possible generators. Lastly, we propose a new tool, called *CIMICE-R*, which implements the described methods, and we evaluate its results on both synthetic and real datasets. *CIMICE-R* has been published as R package on Bioconductor [6].

### Related work

One of the earliest computational models of oncogenesis [7] represented the accumulation of mutations as an *oncogenetic tree* of causal dependencies among alterations, in which the root denotes the wildtype and each path in the tree describes a sequence of causally related events. As in our approach, the nodes in an oncogenetic tree correspond only to observed genetic alterations, with no inferred genotypes. Despite some similarities with our approach, there are two fundamental differences between CPMC and oncogenetic trees: first, CPMC are *Directed Acyclic Graphs* (*DAGs*), not trees; second, and more importantly, the meaning attatched to the edge probabilities is different: in oncogenetic trees, the probability assigned to an edge is the probability of the event “this edge exists”. In CPMC, they instead represent the probabilities of transitioning from one state to the next.

One of the first computational methods to infer an evolutionary mutation tree from SCS data was proposed in [8]. Rather than inferring a phylogenetic tree, their method directly describes temporal ordering relationships among mutations sites by also taking into account sequencing errors. The idea is to compute a “pairwise order relation”, which is a partial temporal ordering on the observed genotypes, represented by a genealogical tree, whose leaves are labeled by the observed genotypes and whose internal nodes correspond to putative common ancestors of the lineages of the samples. Then, mutations are superimposed on the branches of the tree, so that either a mutation temporally precedes another, or two mutations are considered independent. In other words, the ordering is determined by set inclusion: when a genotype has a subset of mutations of another, then the former must temporally precede the latter. For instance, if 00 encodes the wildtype, and 01, 11 are two other observed genotypes (with one and two mutations at the considered sites, respectively) then the inferred temporal ordering is 00 → 01 → 11. To deal with situations that are inconsistent with the above rule, e.g., a triple 01, 10, and 11 of observed genotypes encoding a branching evolution from the wildtype, a Bayesian approach is incorporated into the method. A CPMC provides information similar to the genealogical tree of [8], but since CPMCs are DAGs, branching lineages of evolution can be trivially modeled in the graph structure. The previous “inconsistent” example would be modeled as a CPMC with four edges and a diamond topology: 00 → 01, 00 → 10, 01 → 11, 10 → 11.

OncoNEM [9] is an automated method based on a nested effects model for reconstructing clonal lineage trees from noisy somatic SNV data of single cells. OncoNEM works by clustering together cells with similar profiles; then, it infers their genotypes and unobserved ancestral genotypes; finally, it outputs the inferred tumor subclonal compositions, an evolutionary tree describing the history of such subpopulations, and posterior probabilities of the occurrence of mutations. OncoNEM’s algorithm works by assigning a probabilistic score to sets of mutations and by searching for high-scoring models in the space of possible trees.

SCITE [10] is also a max-likelihood search algorithm that infers the evolutionary history of a tumor from noisy and incomplete SNV data, but, unlike OncoNEM, it focuses on mutation trees. SCITE makes the infinite sites assumption, hence it assumes that the input matrix describes a perfect phylogeny.

An important limitation of both OncoNEM and SCITE is that they work under the infinite sites assumption, i.e., under the hypothesis that each mutation may only occurs once in the evolutionary tree. Evidence has been brought forward to show that real SCS data violates that assumption, and that finite-site models taking into account chromosomal deletions, loss of heterozygosity and convergent evolution lead to more accurate inference of tumor phylogenies [11], [12], [13], [14]. Although our approach assumes that mutations are never lost, we do allow for convergent evolution.

Classic phylogenetic approaches, such as UPGMA and neighbour-joining [15], [16], [17], and other kinds of clustering methods [18], have also been applied to SCS data. Building correct phylogenies with such methods can be done efficiently under the infinite sites assumption if the data contains no errors and mutations persist generation after generation [19]; under less restrictive hypotheses, however, they tend to be outperformed by the more focused approaches described above.

To improve the accuracy of variants detection, single-cell specific variant callers should be used. Monovar [20] and SCcaller [21] were the first two callers developed specifically for SCS data; SCIΦ [22] and SCAN-SNV [23] are two more recent approaches to solve the same problem.

One limitation of our proposal is that it assumes that subclonal reconstruction has already been performed and clonal genotypes have been resolved. Rather than including a specific inference method into our model, we rely on tools such as SiCloneFit [13], Single Cell Genotyper [24] or BEAM [25] to provide the required input.

Although the technology is continually improving, the number and size of published SCS data sets are still limited. A few tools exist that permit generating simulated SCS data sets, and in some cases also inferring their phylogenies [26], [27].

Another way to tackle the lack of high-quality high-throughput SCS data is to develop statistical models that combine such data with traditional bulk sequencing data [28], [29], [30], [31], [32]. In this paper, however, we consider only SCS data.

Finally, the literature on SCS and computational analysis is too large to be summarized exhaustively. Several surveys on various methods and tools for inferring tumor histories from single-cell genomic data have been published to date, including [2], [3], [4], [19], [33], [34], [35], [36], [37], [38], [39], [40], [41].

## Results and Discussion

In this work, we consider a DAG model suggestebd by the following general intuition. Phylogenetic trees put each existent taxon (e.g., a cell from an SCS experiment) in a leaf, and the internal nodes are their inferred extinct ancestors. However, in a tumor ancestral clones coexist with more recent ones, so a phylogenetic tree does not seem the best way to capture the SCS data. DAG models allow to represent more complex evolutionary trajectories and are especially suited for convergent evolutions. In fact, our method is not based on the “infinite sites” assumption, and we do not assume that a perfect phylogeny exists for a set of cells, i.e., the set of cells having a given mutation *m*_1_ and the set of cells having another mutation *m*_2_ may well be properly overlapping.

We identify a minimal set of assumptions on tumor evolution ensuring that DTMCs having DAGs as support correctly model the disease progression. Intuitively, DTMCs are probabilistic models in which the next state of a system only depends on the current one. In our context the state of the system is the genotype of a tumor cell. The tumor cell will generate new cells whose genotypes will represent the next state.

We assume that:

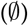 the evolution starts from “normal” cells, i.e., cells which exhibit the same genotype of healthy cells of the same patient;
(∪) mutations can only be acquired along the progression of the disease;
(MC) the probability that a cell will generate cells with new mutations only depend on the genotype of the cell itself;
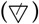 a minimal number of mutations is acquired in each new generation of cells.

DNA SCS data support hypothesis 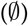 and (∪) since the DNA sequence is not influenced by the cell’s life cycle and all mutations can be detected independently from the gene expression levels.

We are not pretending that these assumptions completely describe the high level of complexity of tumor evolution. Instead, we are trying to reason on the smallest possible set of hypotheses that allows us to rely on DTMCs as modeling formalism and to infer the underlying chain from a dataset. While hypothesis (MC) allows the use of DTMCs, the other hypotheses guarantee that the model has a simple topology, i.e.:

- it is acyclic thanks to (∪);
- it has a single source thanks to 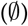;
- it has no “forward” edges allowing to jump intermediate states thanks to 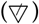.

Agreeing on the above assumptions, we propose a method to infer a DTMC that models the mutational evolution of a tumor from a dataset of genotypes collected from cancer cells. In particular, our method takes in input a Mutational Array containing the genotypes of a set of cells taken from the tumor at a single time and output a DTMC that satisfies the above hypothesis, which we call CPMC. The dataset has to be representative of all the genotypes present in the tumor, i.e., it has to reflect the genotype distribution in the tumor. When in the dataset there are not two possible explanations for a genotype (no convergences) we formally prove that the CPMC that we output is the only model that satisfy our hypothesis. When there are convergences we output one of the possible CPMCs that explains the data.

We do not aim at correcting errors in the input data within our method. For this task we rely instead on other tools capable of clustering and cleaning mutational matrices based on known or predicted false positive, false negative and missing value rates. As an example of such approach, the CIMICE R package provides an easy to use interface to SicloneFIT’s preprocessing algorithm.

To further reduce the dependency of the results from random noise in the data we implemented a bootstrap based approach that consists in the random resampling of the input mutational matrix’s rows. This allows CIMICE to produce many different CPMC models that are finally merged, helping the user to identify nodes and edges that might be generated only because of the noisy nature of SCS data. The merging operation is done naturally by averaging the weighted adjacency matrices of the CPMCs produced by running CIMICE on the different sampled datasets.

To asses the performances of CIMICE, we test its usage on both artificial datasets generated accordingly to our model with different levels of noise and two real world case studies.

As for the simulation, the datasets were generated from the graph in Figure 1, setting the length of the generated path *k* to 5 and simulating 100 cells. We repeated the simulation 4 times for the following false positive *FP* and false negative *FN* rates:

- *FP* = 0.01 and *FN* = 0.05
- *FP* = 0.01 and *FN* = 0.10
- *FP* = 0.01 and *FN* = 0.15

**Fig. 1:**
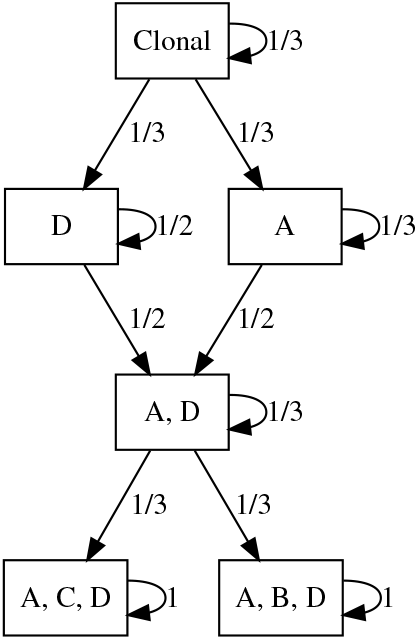
The CPMC used to generate the example datasets. Random paths of fixed length are simulated from the clonal node to generate the genotype of a single cell.

In Figures 2 and 3, we test and compare results between CIMICE and CIMICE copuled with SiCloneFit’s preprocessing algorithm. To produce each result we resampled 100 cells from the generated dataset 1000 times. Other simulations and a comparison with SCITE are reported in the Supplementary Material.

**Fig. 2:**
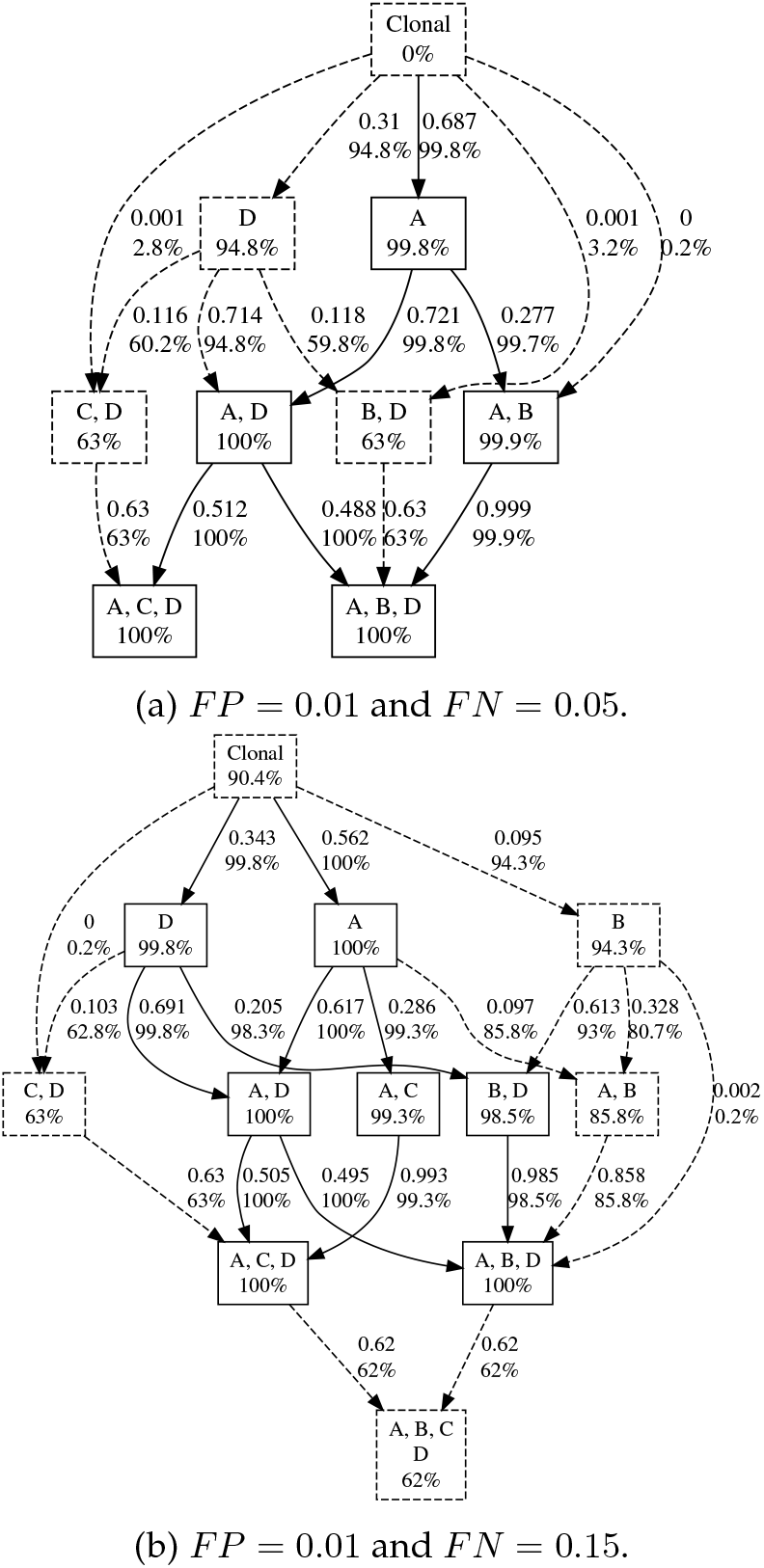
Examples without SiCloneFit preprocessing. Dashed components have a bootstrap probability less than 95%.

**Fig. 3:**
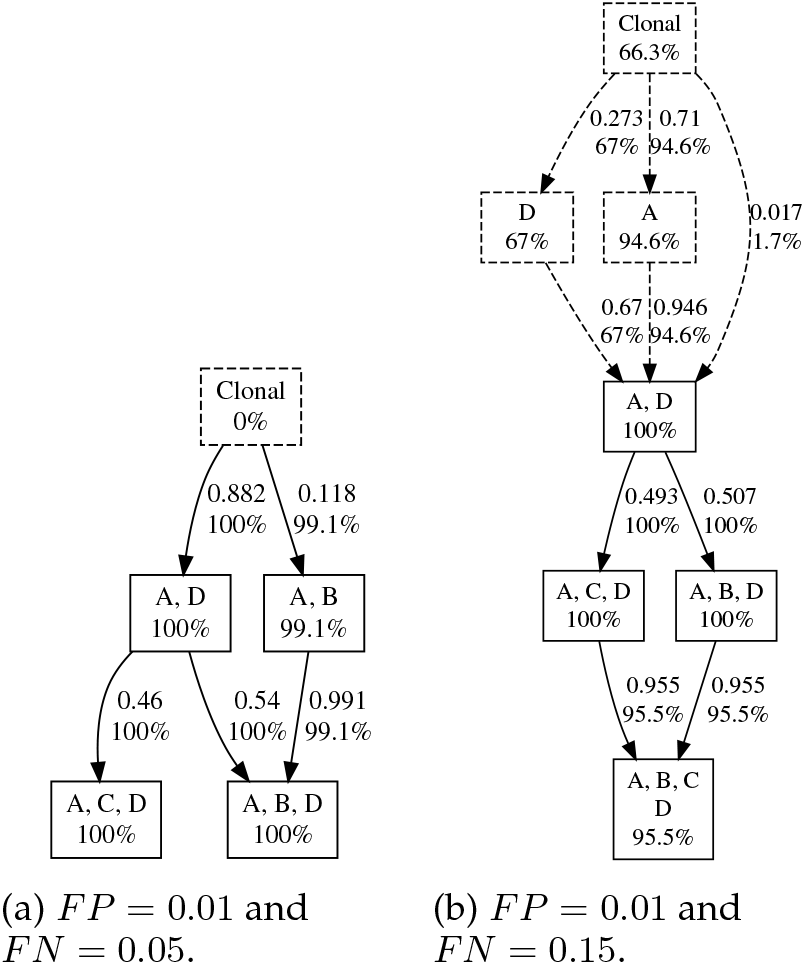
Examples with SiCloneFit preprocessing.

Finally, we test our approach on real datasets, specifically in two settings. In the first we considered a a dataset on clear cell renal cell carcinoma from [17], and in the latter metastatic colorectal cancer data from [42] (CRC1). We show the results of our method coupled with SiCloneFit preprocessing, using the False positive, negative and missing rates reported in the literature (Figures 10 and 11). We set the bootstrapping method to resample 100 datasets with the same size of the original ones.

## Methods

### Setting the Biological and Experimental Context

We represent the state of a cell as the set of mutations present in it. A normal cell is considered to harbor no mutations. Such absence of variants can be defined by exploiting either an external reference or the the healthy cells of the patient. The first method requires attention in mutation selection, while the latter may hide genetic predisposition to tumor development. A cell in any other state than the normal one is possibly dangerous.

Formally, let *G* = {*g*_1_,…, *g_n_*} be a set of *gene mutations*. A set *S* ⊆ *G* denotes the *genotype* over *G* for a cell in which the mutations of *S* are present, while the mutations in *G* \ *S* are not. In particular, 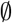 is the genotype of a normal cell. The set of all possible genotypes over *G* is 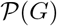, the powerset of *G*.

We are interested in the reconstruction of a probabilistic model representing the mutational history of a cell, i.e., the temporal sequence of the genotypes of the cell’s ancestors. To this aim, some assumptions on the mechanisms underlying the mutational events are needed. We formulate the following **Model’s Hypotheses**:

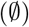 The normal cell 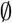 is an ancestor of every cell.
(∪) Mutations can only be acquired, and multiple mutations may be acquired from one cell generation to the next.
(MC) The probability of a mutational event in a cell only depends on its current genotype—that is, it does not depend on how the cell reached a certain state.
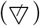 An evolutionary history is anti-transitive and minimal, in the sense that it does not contain another evolutionary history that can explain the same observed genotypes, subject to the requirement that it must account for every plausible trajectory—that is, if *X* ⊂ *Y* are two genotypes then there must be a path from *X* to *Y*.

The *emptyset* hypothesis 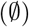 states that each mutational history always starts from a normal cell. This hypothesis can be relaxed without significantly affecting the results in this paper. For instance, if there are some mutations that are present in all the cells of the system under analysis, the 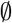 genotype can be replaced by a given genotype containing the mutations acquired at birth by the patient. The *union* hypothesis (∪) specifies that mutations are never lost, i.e., the genotype of an ancestor of the current genotype is a subset of the current genotype. The *homogeneous Markov chain* hypothesis (MC) states that the acquisition of mutations is probabilistic and can be modeled through Markov chains, since each genotype uniquely determines the probability of transitioning to any other genotype. The *anti-transitivity* hypothesis 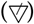 asserts that whenever it is possible to observe a sequence of transitions from a genotype to another, it is not possible to observe any of its subsequences. This is a sort of parsimony assumption, because it implies that each new mutated generation only acquires a minimal number of mutations.

#### Example 1.

*Consider the set of mutations* {*A, B, C, D*} *and a cell having genotype ABC. We are interested in inferring a probabilistic model in which the genotypes of the ancestors of this cell are represented. From the* 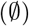 *hypothesis the normal genotype is an ancestor of the cell. The* (∪) *hypothesis rules out the possibility that an ancestor of the cell has genotype ABCD. Without further information, any other genotype* (*A, B, C, AB, AC, and BC*) *may have potentially appeared in the mutational history of a cell with genotype ABC (and different cells with the same genotype may have followed different evolutionary trajectories), although the true history of a specific cell with genotype ABC can involve at most three different ancestor genotypes. If a history contains three ancestor genotypes then either zero or one mutation was acquired, starting from* 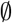, *at each new generation* (*see below*).

*Let us further assume that we know the following:*

1. *normal cells can generate cells with genotype A, but never generate cells with genotype B or C;*
2. *a cell with genotype A can generate cells with genotype either AB or AC.*

*Item (1) does not explicitly forbid that a normal cell generate a cell with genotype, say, AB. However, since a cell with genotype A may be generated from* 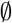, *the* 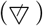 *hypothesis excludes that* 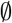 *directly gives rise to an AB cell. Similarly, item (2) together with hypothesis* 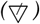 *implies that a cell with genotype A cannot (directly) generate a cell with genotype ABC, because of possible trajectories passing through AB or AC. Therefore, given the additional knowledge the two possible mutational histories of the cell are* 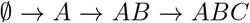 *and* 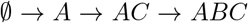. *Note that each arrow corresponds to a mutated new cell generation: but it is possible that the same genotype is maintained for many generations. Finally, by hypothesis (MC), each arrow can be labelled with a transition probability, giving rise to a graph probabilistic model (Figure 4). Such probabilities are determined from experimental data as detailed below.*

**Fig. 4:**
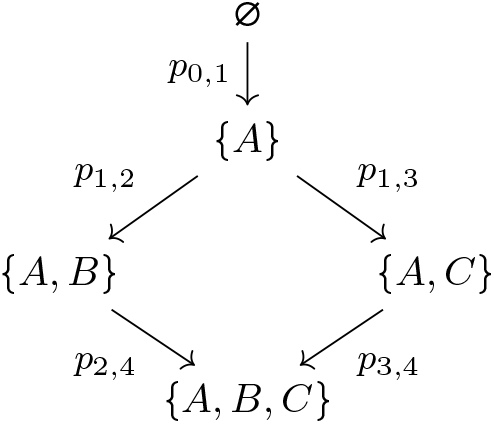
A generic probabilistic model for Example 1

We focus on SCS data and in particular on DNAseq data, as the DNA molecule offers chemical stability properties that support our hypotheses. Moreover, the DNA sequence is not influenced by the cell’s life cycle and all mutations can be detected independently from the gene expression levels.

Aware of the limitations and errors of the current SCS technologies, in this work we consider an ideal setting in which all the relevant mutations are correctly detected and a large number of cells from a tumor region is analysed. As the technology improves, it is reasonable to assume that larger datasets will become available and that the rate of errors will decrease. Currently, to approximate this ideal setting the data may be preprocessed with tools that impute missing values and resolve clonal genotypes [13], [24], [25]. A possibility to derive relatively large datasets is to preprocess data from bulk sequencing experiments and extract plausible single cell explanations [43].

As we will see, working on SCS data has the following advantages with respect to bulk sequencing data:

- we can drastically simplify the model inference engine, since the set of genotypes present in the tumor are represented in the data and need not to be inferred;
- we can formally prove that when each cell has a unique possible set of ancestors, the produced model is *correct*, i.e., no other information is needed;
- we propose models that can be used to generate artificial data.

A SCS experiment consists in the sequencing of a set of cells taken from either *in vivo* or *in vitro* samples. Hence, the genotype of each analysed cell is known. Usually, the results of such experiments are represented through Boolean matrices, called *Mutational Arrays* [3], in which each row represents a cell and each column represents a mutation. The value in position (*i,j*) of a mutational array is 1 if and only if the *j*-th mutation is present in the *i*-th cell. Mutational arrays have a broad usage among many tools in the field of tumor phylogenetics (see, e.g., [44], [45]).

As for the underlying models, we need some assumptions on the data as well. In particular our **Data Hypotheses** are:

(ONE) All the analysed cells are taken at the same time from a single site, i.e., they represent *one* snapshot of a cancer tissue.
(POP) The analysed cells reflect the genotype distribution of the *population* of all the cells in a given site.

Under these assumptions, given a mutational array it makes sense to define a frequency distribution over the genotypes.

#### Definition 1

(Dataset Distribution). *Let G be a set of gene mutations. A* dataset distribution *D* over *G is a frequency distribution over the genotypes of G, i.e., a function* 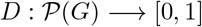 *with* 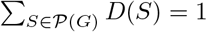.

In what follows, we will omit the underlying mutational arrays and refer to the corresponding dataset distributions, called simply *datasets* hereafter. A dataset is typically defined by its support: although the size of 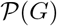 is exponential with respect to the size of *G*, usually only a limited number of genotypes is observed, so the support usually has a small size.

#### Example 2.

*Consider again the set of genes* {*A, B, C, D*}. *A possible mutational array over ten cells, and its corresponding dataset distribution, is shown below*:

| | $A$ | $B$ | $C$ | $D$ |
| --- | --- | --- | --- | --- |
| $c_1$ | 0 | 0 | 0 | 0 |
| $c_2$ | 1 | 0 | 0 | 0 |
| $c_3$ | 1 | 0 | 0 | 0 |
| $c_4$ | 1 | 1 | 0 | 0 |
| $c_5$ | 1 | 0 | 1 | 0 |
| $c_6$ | 1 | 0 | 1 | 0 |
| $c_7$ | 1 | 1 | 1 | 0 |
| $c_8$ | 1 | 0 | 0 | 0 |
| $c_9$ | 1 | 0 | 1 | 0 |
| $c_{10}$ | 1 | 1 | 1 | 0 |

*Dataset distribution*:

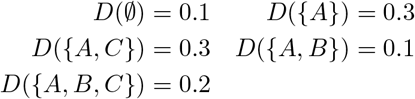

Our goal is to find a plausible probabilistic model of the mutational histories of the cells in a given dataset, within a certain class of Markov models based on the previously stated assumptions. In general, such a problem does not have a unique solution, even in an ideal generalized setting in which an infinite sequence of datasets corresponding to temporal snapshots of a sequenced tissue is available (see Example 7). In order to overcome such difficulty, we will (a) identify a few additional conditions guaranteeing the uniqueness of the model reconstructed from a dataset, and when uniqueness cannot be achieved, explicitly describe what missing information prevents that; and (b) when such additional information is not available, propose a reasonable criterion for the reconstruction of an admissible model.

### Basics on Discrete Time Markov Chains

Given our model’s hypotheses, it is reasonable to consider DTMCs as the underlying mechanism generating the data. The nodes of such chains correspond to the possible genotypes of a cell, while the edges model the probability of a genotype to mutate, i.e., to acquire new mutations. In this section we briefly report some basic definitions and introduce the notations we use on DTMCs. We refer the reader to [46], [47] for a complete presentation on the topic.

#### Definition 2

(Discrete Time Markov Chain (DTMC)). *A* Discrete Time Markov Chain (DTMC) *is a pair M* = (*V,p*), *where*:

1. *V is a finite set of vertices;*
2. *p*: *V* × *V* → [0,1];
3. ∑_*v*∈*V*_ *p*(*u, v*) = 1 *for each u* ∈ *V*.

Usually, when a DTMC is considered an initial probability distribution on the states of the chain is also given. The initial distribution specifies the probability of being in each state of the chain at time 0. So, from the initial distribution and the edges probabilities, one can compute the probability distribution over the states of the chain at time *t*, i.e., after crossing *t* edges. We anticipate here that in our context at time 0 the normal genotype has probability 1 and given the distribution at time *k* we try to infer the probabilities of the edges.

A DTMC can be represented as a graph whose edges are labeled with probabilities, where an edge is present only if its probability is greater than 0. For each vertex of the graph the sum of the probabilities over its outgoing edges is 1. We adopt the following standard graph notation:

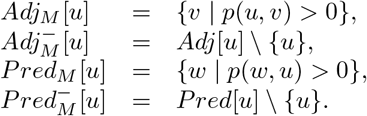

The subscript *M* may be omitted when the chain is clear from the context.

In DTMCs the quantity *p*(*u, v*) that labels the edge (*u, v*) represents the probability that the chain is in *v* at time *t* + 1, given that it was in *u* at time *t*, i.e., *p*(*u, v*) = *P*[*X*(*t* + 1) = *v* | *X*(*t*) = *u*]. Similarly, we can consider the probability that the chain was in *u* at time *t*, given that it is in *v* at time *t* + 1, i.e., *P* [*X* (*t*) = *u* | *X* (*t* + 1) = *v*]. Note that, while *P*[*X*(*t* + 1) = *v* | *X*(*t*) = *u*] = *p*(*u, v*) does not depend on *t* by definition of DTMCs, in general *P*[*X*(*t*) = *u* | *X*(*t* + 1) = *v*] does depend on *t*.

We say that a vertex *u reaches* a vertex *v* if there exists a finite sequence *u*_0_, *u*_1_,… *u_m_*, called a *path*, such that *u*_0_ = *u, u_m_* = *v*, and *u*_*i*+1_ ∈ *Adj*[*u_i_*] for each *i* ∈ [0, *m* – 1]. The probability of the path is the product of the probabilities of the edges along the path. The probability of reaching *v* from *u* is the sum of the probabilities of the paths from *u* to *v*. A *cycle* is a path from a vertex *u* to itself involving at least another vertex different from *u*, i.e., we do not consider self-loops as cycles. We say that a DTMC is *acyclic* if it has no cycles (an acyclic DTMC may have self-loops).

As already mentioned, in a DTMC the probability *p*(*u, v*) represents the probability of moving from *u* to *v*, when a tick of the clock occurs, i.e., time moves from *t* to *t* + 1. Hence, we are considering an underlying global time that flows homogeneously with respect to the state of the system. On the other hand, we could be interested in a model where the clock ticks only when there is a change of state, i.e., the ticks of the clock measure the number of jumps from one state to another. In DTMC literature, this is known as the *jump* version of a chain.^1^ The jump version of a chain can be computed by assigning to each edge (*u, v*) with *u* ≠ *v* a probability proportional to *p*(*u, v*) in such a way that the sum of the probabilities of the edges from *u* to *V* \ {*u*} is 1. When *Adj*[*u*] = {*u*}, the self-loop is assigned probability 1.

#### Definition 3

(Jump DTMC associated to a DTMC). *Let M* = (*V, p*) *be a DTMC. The* Jump DTMC associated to *M is the DTMC J*(*M*) = (*V, jp*), *where, for every u, v* ∈ *V, jp*(*u, v*) *is defined as follows*:

- *if Adj_M_* [*u*] = {*u*}, *then jp*(*u, u*) = 1 *and jp*(*u, v*) = 0 *for every v* ≠ *u*;
- *otherwise, jp*(*u, u*) = 0 *and*

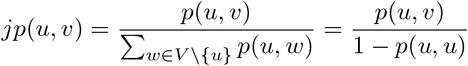

*for every v* ≠ *u*.

### Cancer Progression Markov Chains

In our context we refer to a subset of DTMCs that we call Cancer Progression Markov Chains (CPMCs). CPMCs have additional properties which make them admissible as models of cancer progression. Intuitively, the vertexes of a CPMC represent the genotypes involved in the cancer progression under analysis. The empty genotype is the normal one that is at the origin of every mutational history. Every other vertex is reachable from the empty genotype. Since, under our hypotheses, mutations cannot be removed, a vertex representing a genotype cannot reach another vertex representing a genotype with less mutations. As a consequence, CPMCs are always acyclic. Moreover, since we are assuming that the evolutionary history is always the one involving less mutations, whenever there is a path of length at least 2 from one vertex to another, there cannot be an edge connecting the two vertices. This implies that CPMCs are anti-transitive. The above observations lead to the formulation of the following definition of CPMCs.

#### Definition 4

(Cancer Progression Markov Chain (CPMC)). *Consider a set of genes G* = {*g*_1_,…,*g_n_*} *and let* 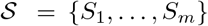 *be a set of genotypes over G, with* 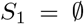. *A* Cancer Progression Markov Chain 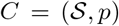 *over* 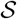 *is a DTMC such that*:

1. 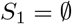 *reaches any other genotype of the chain;*
2. *for every i, j* ∈ [1, *m*], *p*(*S_i_, S_j_*) > 0 *if and only if S_i_* ⊆ *S_j_ and there is no k* ≠ *i, j such that S_i_* ⊆ *S_k_* ⊆ *S_j_*.

The first condition of our definition of CPMC states that the normal genotype 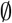 is always present and it is the initial state of any mutational evolution. In other terms, in CPMCs we are always implicitly considering the initial distribution that at time 0 gives probability 1 to 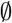 and 0 to all the other states.

In the second condition of the above definition we have been more restrictive than stated in our hypotheses. In particular, we have imposed that whenever a genotype *S_i_* is one of the minimal explanations for a genotype *S_j_*, the probability of going from *S_i_* to *S_j_* is greater than 0. This restriction is not too demanding, since such probability can be arbitrarily small. It allows us to uniquely define the topology (i.e., the set of edges) of the chain for a given set of genotypes. However, it is possible to drop such restriction when further information on the topology is available.

#### Example 3.

*Let us consider the set of genes G* = {*A, B, C*} *and the set of genotypes* 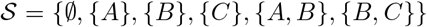. *In Figure 5 we represent two possible CPMCs over* 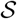.

**Fig. 5:**
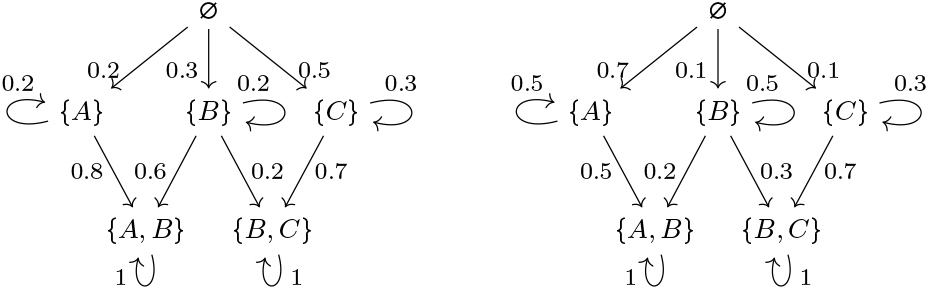
CPMCs over 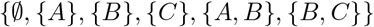.

Since a CPMC is a DTMC, given a CPMC *C* we can build the jump DTMC *J*(*C*) associated to *C*. The properties of *C* ensure that also *J*(*C*) is acyclic, with a single source vertex, and anti-transitive. In particular, *J*(*C*) is still a CPMC.

#### Lemma 1.

*Let C be a CPMC. Then is acyclic, anti-transitive, and J*(*C*) *is a CPMC.*

#### Example 4.

*Let us consider the two CPMCs depicted in Figure 5. Figure 6 represents their associated jump chains.*

**Fig. 6:**
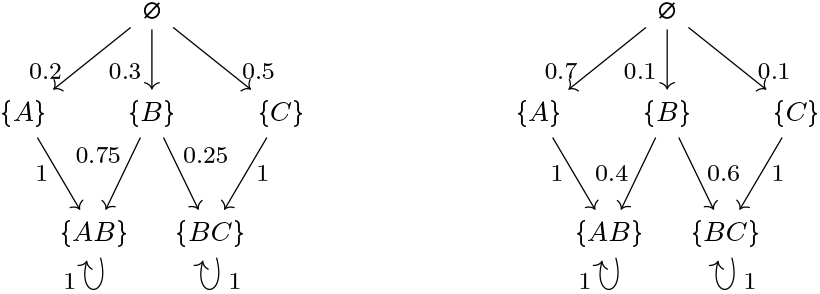
Jump chains. The jump chains associated with the CPMCs depicted in 5.

### Cancer Progression Markov Chains and Datasets

Let us assume that we know that the evolution of a type of cancer is regulated by a given CPMC *C*. We can use *C* to generate simulations of the evolution of the cancer. Moreover, we can use *C* to determine the probability that a cell with a given genotype will degenerate into another one.

Notice that in CPMCs time evolves, i.e., edges are crossed, when a cell cycle is completed. However, it makes no difference in our context to replace single cell cycles with their multiples, e.g., consider the new state after 100 cell cycles, or even with periodic observations of the system. On the other hand, we could have referred to Continuous Time Markov models in which time can be expressed in days, months, years (depending on the desired granularity). In that case probabilities would have been replaced by transition rates. However, without more specific knowledge on proliferation/death rates of different genotypes, continuous time models would give us an equivalent view.

Interestingly CPMCs can be used as data generators to validate other inference methods, provided that such methods agree on our four Model Hypothesis. In particular, we can randomly generate a CPMC *C*, use it to generate a dataset *D_k_*, where *D_k_* (*S*) is the probability that *C* is in state *S* at time *k*, i.e., *D_k_* (*S*) = *P*[*X*(*k*) = *S*], apply the inference method on *D_k_* and check whether the inferred knowledge is correct with respect to the underlying chain *C*. This process can be repeated until we are able to either accept or reject the inference method. The CPMCs can also be artificially engineered in order to test the behavior of the method on limit cases.

We recall that, since 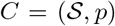 is a Markov Chain and we assume that at time 0 the process starts from the normal genotype 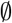, we have:

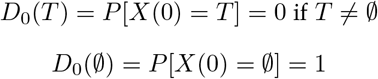

and

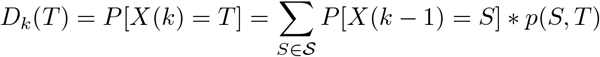

for each *k* > 0

#### Example 5.

*Let us consider again the CPMC C depicted in Figure 5 on the left. The dataset D*_0_ *generated by C at time* 0 *is* 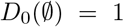. *The dataset D*_1_ *generated by C at time* 1 *is D*_1_({*A*}) = 0.2, *D*_1_({*B*}) = 0.3, *and D*_1_({*C*}) = 0.5. *The dataset D*_2_ *generated by C at time* 2 *is D*_2_({*A*}) = 0.2^2^, *D*_2_({*B*}) = 0.3 * 0.2, *D*_2_({*C*}) = 0.5 * 0.3, *D*_2_({*A, B*}) = 0.2 * 0.8 + 0.3 * 0.6, *and D*_2_({*B, C*}) = 0.3 * 0.2 + 0.5 * 0.7. *The dataset D*_3_ *generated by C at time* 3 *is D*_3_({*A*}) = 0.008, *D*_3_({*B*}) = 0.012, *D*_3_({*C*}) = 0.045, *D*_3_({*A, B*}) = 0.408, *and D*_3_({*B,C*}) = 0.527.

A given dataset may be generated by different CPMCs at different times (Example 6). Besides, different CPMCs can even generate the same (infinite) sequence of datasets (Example 7).

#### Example 6.

*Let us consider the two CPMCs depicted in Figure 7. Let C*_1_ *be the chain on the left and C*_2_ *be the one on the right. It is immediate to observe that the dataset* 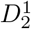 *generated by C*_1_ *at time* 2 *is* 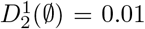 *and* 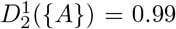. *Such dataset coincides with the dataset* 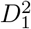 *generated by C*_2_ *at time* 1.

**Fig. 7:**
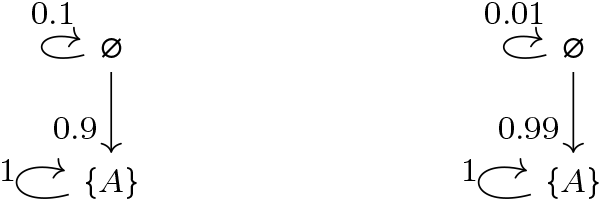
Two CPMCs that generate the same datasets at different time instants.

*We will come back on this example in the next section. The problem here lies in the inference of the probability of the self-loop on* 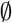. *As a matter of fact both C*_1_ *and C*_2_ *have the same jump chain and we will prove that such jump chain can be inferred exactly.*

When inferring a CPMC from a single dataset *D*, it may not be possible to accurately estimate the time at which the snapshot was taken. The example above shows that, in general, *D* may be supported by different CPMCs, which generate *D* at different times.

Unfortunately, in the worst case two different CPMCs can generate the same datasets at each time instant (Example 7). This is not a problem when CPMCs are used as data generators because the CPMC is known, it for inference it means that in general uniqueness of the model cannot be guaranteed. In the next section we will prove that this can happen only in presence of convergences, i.e., when genotypes have many possible ancestors. In that case we will provide a heuristic which allows us to infer one of the possible underlying jump chains.

#### Example 7.

*Let us consider the two CPMCs depicted in Figure 8. They generate the same datasets at each time instants. As a matter of fact, the first* 3 *levels of the chains are equal with equiprobable branching, while at the last level for each node the sum of the probabilities of the incoming edges is the same in the two chains. The dataset D*_3_ *at time* 3 *for both chains is D*_3_({*A, B, D*}) = 0.3 * 0.7, *D*_3_({*A, C, D*}) = 0.3 * 1.1, *and D*_3_({*B, C, D*}) = 0.3 * 1.2.

**Fig. 8:**
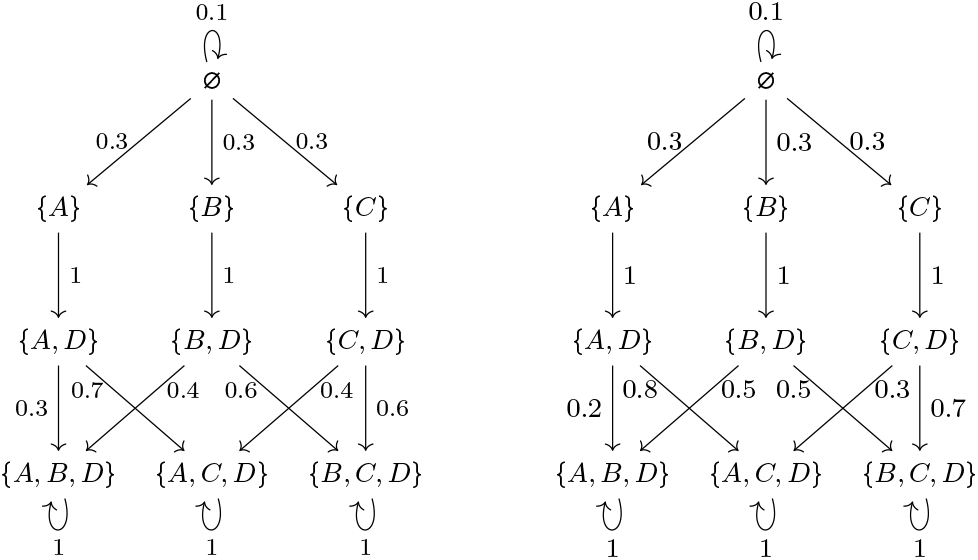
Two CPMCs that generate the same datasets at each time instant.

### Properties of Cancer Progression Markov Chains

Let *f_i_*(*S*) be the event *X*(*i*) = *S*∧*X*(*i*–1) ≠ *S*∧⋯∧*X*(0) ≠ *S*, i.e., the chain is in state *S* at time *i* and has never been in *S* before. Hence, *P*[*f_i_*(*S*)] is the probability of *f_i_*(*S*), i.e., the probability of reaching for the first time the vertex *S* after *i* steps. This is also known as the *first passage probability*. The probability of being in state *T* ∈ *Adj*[*S*]^-^ for the first time after at most *k* steps, passing through the edge from *S* to *T*, can be expressed as

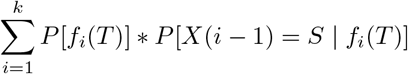

Since *C* is acyclic this is equivalent to

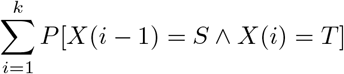

and can be computed on CPMCs as stated by the following lemma.

#### Lemma 2.

*Let* 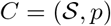 *be a CPMC and let* 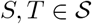 *be such that S* ≠ *T and p*(*S, T*) > 0. *Then*:

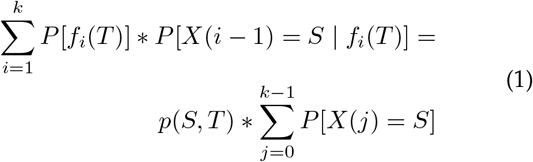

*Proof.* Since *S* ≠ *T and p*(*S,T*) > 0, the edge (*S,T*) is the only path of length 1 from *S* to *T*. We have that:

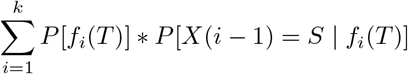

By the definition of conditional probability

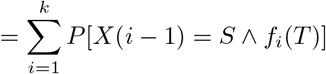

By def. of *f_i_*(*T*), since *C* is a DTMC and (*S, T*) is the only path from *S* to *T*

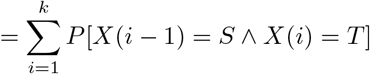

By the definition of conditional probability

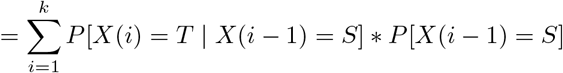

Since *C* is a DTMC

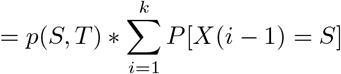

By replacing *i* – 1 with *j*

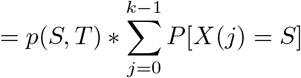

As a consequence we get an alternative method to compute the probabilities on the jump chain *J*(*C*). Let *height*(*C*) be the length of the longest path in *C*, without crossing self-loops. Since *C* is acyclic and the normal genotype 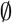 reaches any other genotype, *height*(*C*) is the length of the longest path which does not uses self-loops from 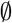 to a leaf in *C*.

#### Theorem 1.

*Let* 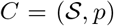 *be a CPMC and k* ≥ *height*(*C*). *Let* 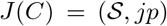 *be the jump DTMC associated to C. Let* 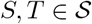 *with S* ≠ *T and p*(*S, T*) > 0.

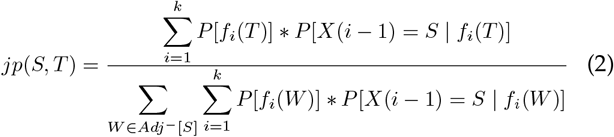

*Proof.* The fact that *k* ≥ *height*(*C*) ensures that the fraction on the right hand side is properly defined, i.e., its denominator is different from 0.

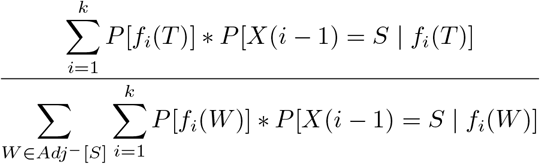

By Equation (1)

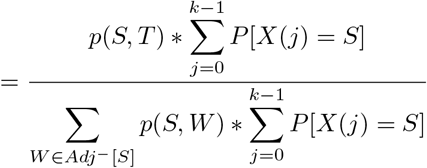

By simplifying

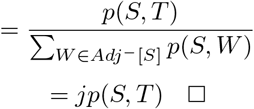

Notice that in Equation (2) the denominator is just a normalization factor which ensures that the sum of the probabilities of the edges from *S* is 1.

### The Inference Method

Let *D_k_* be a dataset satisfying our data hypotheses, representing a snapshot of a tumor after *k* evolution steps. As discussed before, the number of steps in our context models the time elapsed from normality to the observed snapshot. We will see that in our method we do not assume to know the value of *k*. Assuming that *D_k_* has been generated by a CPMC (i.e., by a model satisfying our model hypotheses) one may wonder whether it is possible to infer such a CPMC, i.e., a CPMC such that *D_k_*(*S*) = *P*[*X*(*k*) = *S*] for each genotype *S*, where *k* is not known a priori. To be more precise, since at this point of the construction, we do not want to add information to the dataset, we can say that we are interested in inferring the part of *C* that is visible from the dataset, i.e., the genotypes of *C* will be the normal genotype and the ones having positive frequency in *D_k_*. Formally this means that 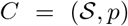, where 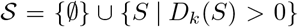. If during the evolution there had been genotypes that have disappeared and are not represented in the dataset our method will not infer such genotypes, since it is our aim to reconstruct a model able to represent the current situation without introducing unobserved knowledge.

Lemma 2 provides a way to compute *p*(*S, T*), but only when 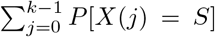 is known. Unfortunately, there is no way to determine such a quantity from the dataset alone. On the other hand, if we consider *J*(*C*) instead of *C* then Theorem 1 can be used to compute the transition probabilities—in some cases exactly, in general using some heuristics.

By definition, the topology of *J*(*C*) is uniquely determined by the set of observed genotypes, as follows:

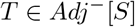

if and only if

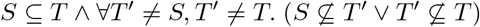

According to Theorem 1, in order to infer *jp*(*S, T*) the following probabilities must be estimated:

a. for each 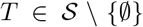 and for each *i* ∈ [1,*k*], the probability *P*[*f_i_*(*T*)];
b. for each 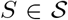, for each *T* ∈ *Adj*^-^[*S*], and for each *i* ∈ [1, *k*], the probability *P*[*X*(*i* – 1) = *S* |*f_i_*(*T*)].

We say that there is a *convergence* in *C* whenever a genotype *T* has two different predecessors, that is, when there is a genotype *T* such that |*Pred*^-^[*T*]| > 1. We distinguish two cases:

1. *C* (or equivalently, *J*(*C*)) has no convergences;
2. *C* (or equivalently, *J*(*C*)) has at least a convergence.

#### No Convergences

If *C* has no convergences, then for each 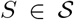, for each *T* ∈ *Adj*^-^[*S*], and for each *i* ∈ [1, *k*] it holds that *P*[*X*(*i* – 1) = *S* |*f_i_*(*T*)] = 1. This is trivial since *S* is the only predecessor of *T*. Hence, by Theorem 1 we get

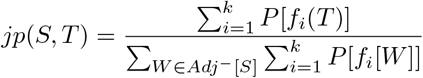

Since the denominator is just a normalization factor, we have to find a way to compute the numerator from *D_k_*.

We can proceed by induction from the leaves to the root of *C*:

- if *T* is a “leaf” of *C*, i.e., 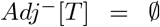, then 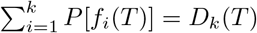;
- otherwise 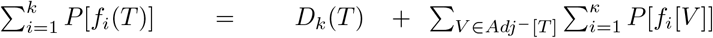.

As a consequence, we have proved the following corollary.

#### Corollary 1.

*Let* 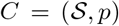 *be an unknown CPMC without convergences and let D_k_ be a dataset generated from C at time k. The chain J*(*C*) *can be uniquely inferred from D_k_, provided that all the genotypes of C are represented in D_k_*.

In other terms, the fact that all the genotypes have to be represented in *D_k_* means that the time instant at which the data are taken is neither too early, so that some genotypes have not yet been discovered, nor too late, so that some genotypes are no more present. Notice that we do not assume to know the value of *k*.

#### Convergences

From the above discussion, it emerges that in the case with convergences we have to find a way to estimate *P*[*X*(*i* – 1) = *S* |*f_i_*(*T*)], for each *i* ∈ [1, *k*]. This means that for any *i* ∈ [1, *k*] we have to estimate the probability that since we are for the first time in *T* at time *i* we were in *S* at time *i* – 1. As already stated in the previous sections, Markov Chains are time homogeneous, but this is not in true in the general case for their reverse. So it is possible that *P*[*X*(*i* – 1) = *S* | *f_i_*(*T*)] ≠ *P*[*X*(*j* – 1) = *S* | *f_j_*(*T*)], for some *i,j* ∈ [1, *k*]. However, without any additional knowledge, the best one can do is approximate such values. In the following, we will approximate all the values uniformly with a single quantity denoted *Split*(*S, T*). In this way, by Theorem 1 we get

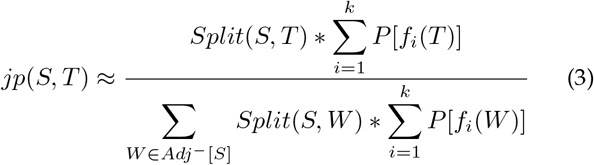

Again, the denominator is a normalization factor, so we focus on the numerator.

- *Split*(*S, T*) is an approximation that we attribute to all the possible values of *P*[*X*(*i* – 1) = *S* |*f_i_*(*T*)] and has to be computed by exploiting only *D_k_*.
- 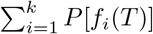 has to be computed by induction from the leaves to the root, but some more caution will be necessary with respect to the case without convergences.

We have already showed that no unique solution may exist in the presence of convergences, i.e., the function *Split*(*S, T*) is not uniquely determined in general (see Figure 8 and Example 7). Our heuristic for *Split*(*S, T*) is based on the following simple considerations.

1. Since *Split*(*S, T*) represents the probability of reaching *T* through *S*, ∑_*X*∈*Pred*^-^[*T*]_ *Split*(*X,T*) = 1.
2. For *S, S*′ ∈ *Pred*^-^ [*T*], if *S* is more frequent than *S*′ in the dataset, then it is more likely that *T* is reached from S than from *S*′.
3. To be more precise, in the previous item not only the frequencies of *S* and *S*′ have to be taken into account, but also those of their ancestors.
4. Also the number of outgoing edges from *S* and *S*′ must be taken into account. If *S* has many outgoing edges, but *S*′ reaches only *T*, then, intuitively, even if *S* and *S*′ have the same frequency, the probability of reaching *T* from *S* should be lower compared to the probability of reaching *T* from *S*′.

Based on the above, we elaborate the following iterative definition for 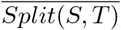, that will be then normalized to obtain *Split*(*S, T*):

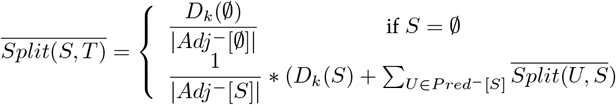

Intuitively, *S* is assigned a weight proportional to its frequency in the dataset and, recursively, to the weight of its ancestors; then, such weight is uniformly distributed over *S*’s outgoing edges. In principle, such distribution should be proportional to *jp*(*S, T*), but since we are still in the process of evaluating it we apply a uniform distribution.

Once all the 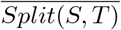 have been computed we can normalize them, thus obtaining the values for *Split*(*S, T*).

In order to compute the values 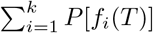 we proceed iteratively from the leaves to the root. However, since a node can have many parents, we cannot assign all its probability to every parent. We use again the heuristic Split to distribute such probability among all parents. In particular, we have:

- if *T* is a “leaf” of *C*, i.e., 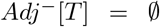, then 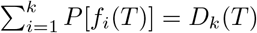;
- otherwise 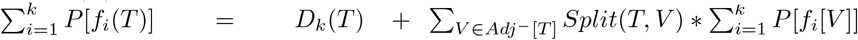.

Finally, we can exploit Equation (3) to get the probabilities *jp*(*S, T*).

If the above heuristic is applied to a topology with no convergences, then all the *Split*(*S, T*) are 1, hence the heuristic computes the same jump probabilities that can be obtained by applying the method described in the previous section for the special case without convergences.

#### Example 8.

*By applying Equation (3) to the dataset D*_3_ *of Example 7 we obtain the jump CPMC depicted in Figure 9. This is the approximation we compute for the jump chains of the models in Figure 8. We recall that both models in Figure 8 are plausible generators for the dataset. Notice that despite the high symmetry of the dataset over the first 7 genotypes, the chain we extract is not completely symmetric. However, we are not inferring the self-loops. It is possible to define a CPMC with self-loop whose jump chain is that presented in Figure 9 and that generates the dataset D*_3_ *by solving a system of equations whose unknowns are the probabilities of the self-loops.*

**Fig. 9:**
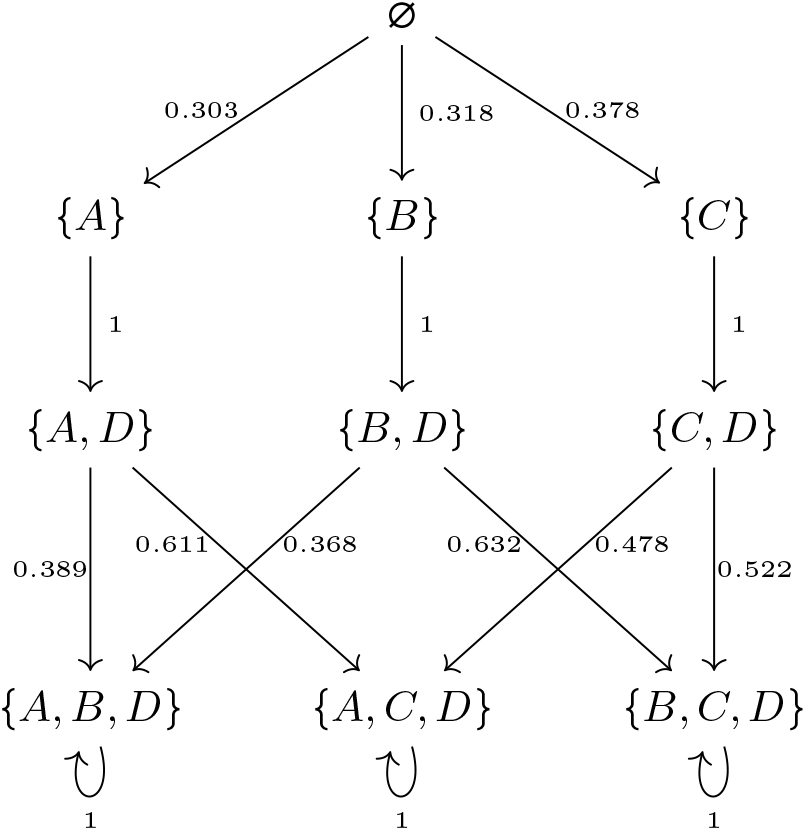
The jump CPMC inferred from the dataset *D*_3_ of Example 7

**Fig. 10:**
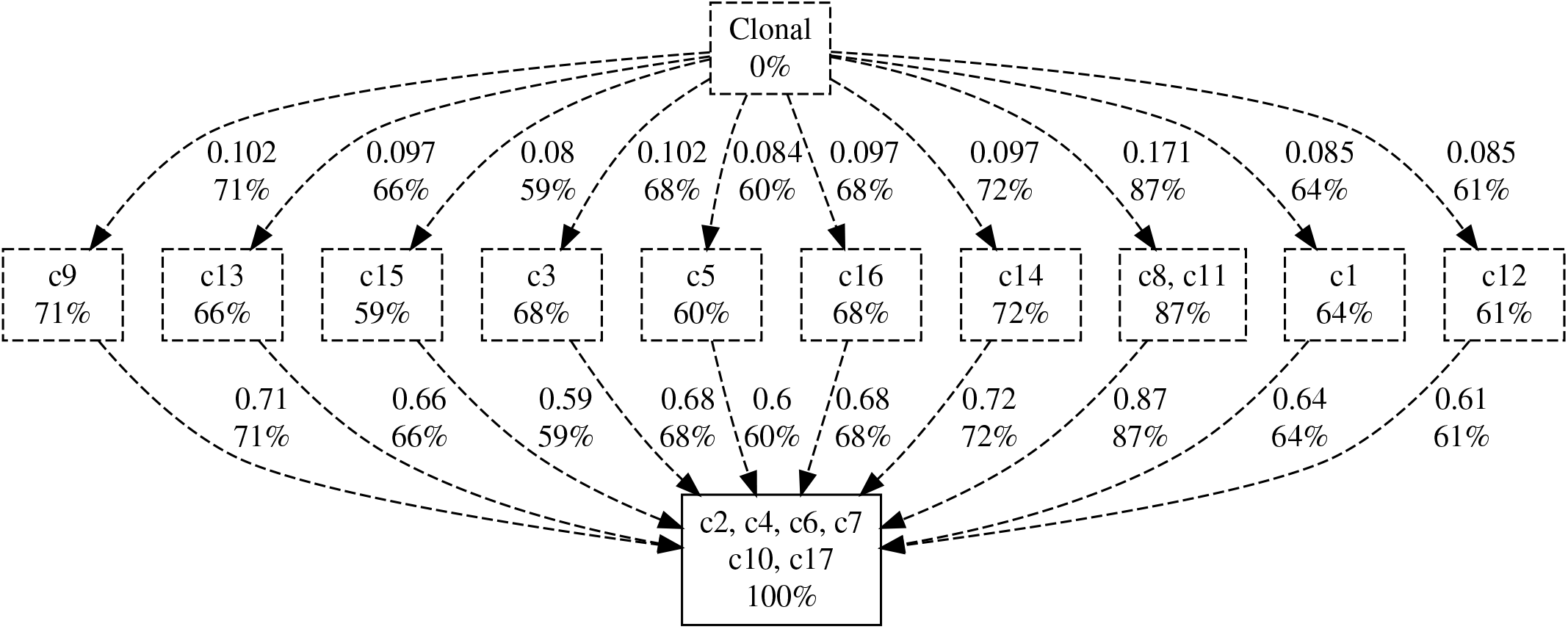
Our method’s results on the dataset from [17]. The False positive, False Negative and Missing Value Rates are 2.67 × 10^-5^, 0.1643, and 0.2117 respectively. Two different stages are clearly separated even if the scarcity and noisiness of data does not allow our method to establish a preferred progression.

**Fig. 11:**
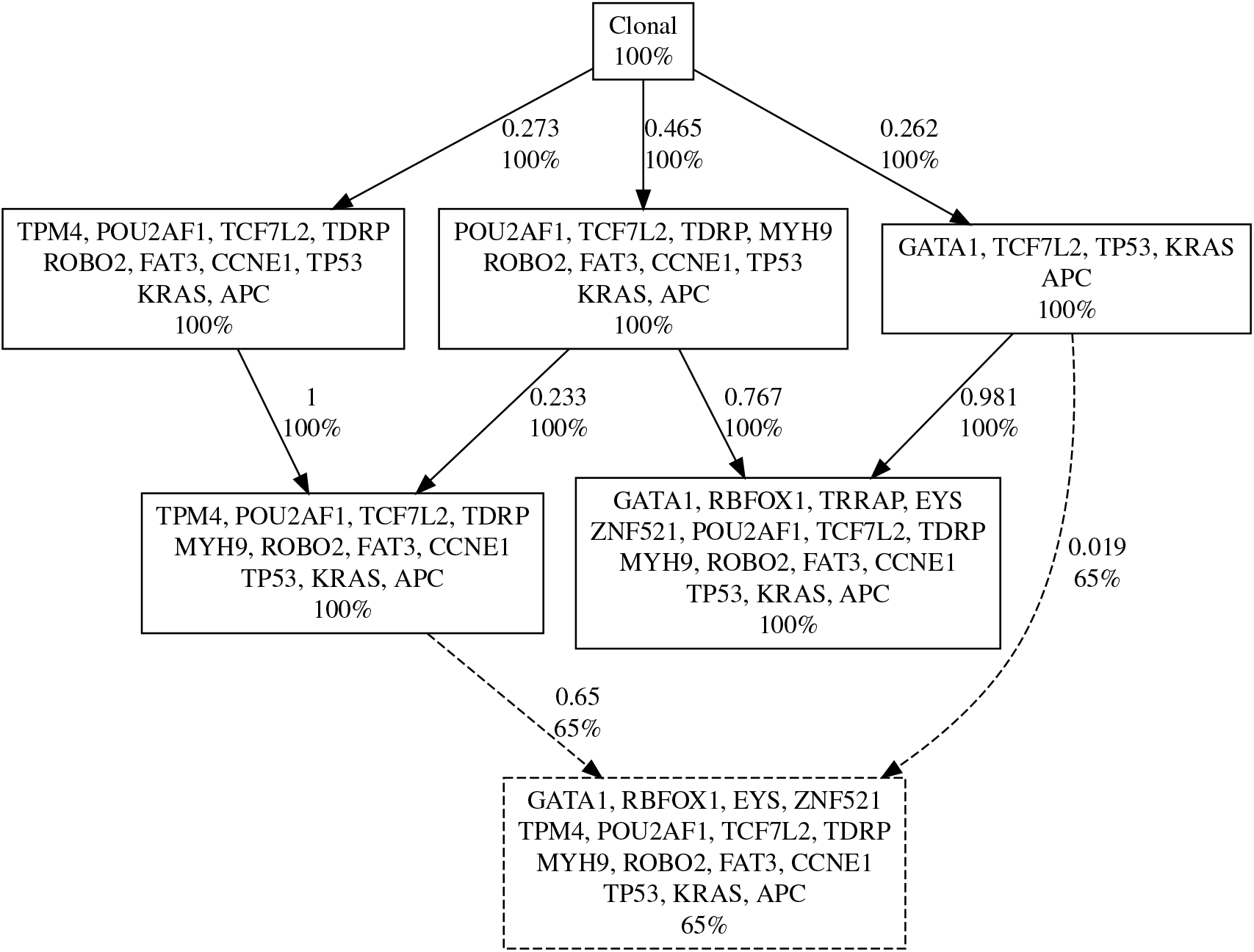
Our method’s results on the dataset from [42] (CRC1). The False positive, False Negative and Missing Value Rates are 0.0152, 0.0789, and 0.0671 respectively. In this case our method reconstruct different progression trajectories, that are mostly in agreement with the subdivision between genes present in metastatic and non metastatic gene assets given in the original paper.

#### All Together

In order to prove the correctness of our method we described it assuming that the dataset *D_k_* has been generated from a CPMC *C*. We demonstrated when and with which accuracy we are able to infer *J*(*C*) from *D_k_*. Summing up we proved the following results.

1. When there are no convergences, we exactly infer *J*(*C*).
2. When there are convergences, if the probability of reaching *T* from *S* is time homogenous and has been estimated, e.g., using further data and experts knowledge, then we can exactly infer *J*(*C*). Notice that such further information is necessary only for the nodes with convergences.
3. When there are convergences and no further information is available, we provided a heuristic for inferring a plausible *J*(*C*).

### *CIMICE-R*: (Markov) Chain Method to Infer Cancer Evolution

The R package *CIMICE-R* implements the above described methods. It takes in input a dataset in form of a mutational matrix, i.e., a boolean matrix representing altered genes in a collection of samples obtained with SCS DNA analysis.

*CIMICE-R* data processing and analysis can be divided in four section: input management, preliminary analysis of the dataset, graph topology reconstruction, chain weight computation, output presentation.

The tool requires a boolean dataframe as input in which each column represents a gene, each row represents a sample (or a genotype), and each 0/1 represents whether a given gene is mutated in a given sample. It is possible to load this information from a file. The default input format is the “CAPRI/CAPRESE” TRONCO [48] format: the file is a tab or space separated file; the first line starts with the string “s/g” (or any other word) followed by the list of genes (or loci) to be considered in the analysis. Each subsequent line starts with a sample identifier string, followed by the bit set representing its genotype. Another option is to define directly the data frame in R. In the case of data composed by samples with associated frequencies it is possible to use an alternative format that we call *CAPRIpop*, where the frequency column is mandatory and samples cannot be repeated. Another option is to compute a mutational matrix directly from a MAF file.

The tool includes simple functions to quickly analyze the distributions of mutations among genes and samples. Correlation plots are also available. In case of huge dataset, it could be necessary to focus only on a subset of the input samples or genes. *CIMICE-R* provides an easy way to do so when the goal is to use the most (or least) mutated samples and/or genes.

The subsequent stage goal is to obtain the topology for the final Cancer Progression Markov Chain. Once the topology has been computed, it can be plotted, e.g., using igraph. Finally, the probabilities that labels the edge of the jump chain are computed. The tool first computes the 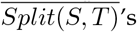. These are called UP weights in the implementation. Then, these are normalized to obtain the *Split*(*S, T*)’s (called normalized UP weights). From these the probabilities can be derived (also called normalized DOWN weights).

In order to show the results of the analysis exploiting different libraries three output methods are provided. These libraries improve on the default igraph output visualization.

All the details and examples of usage are provided at [6].

## Acknowledgment

The authors are partially supported by PRIN 2020 NiRvAna code 20202FCJMH and INdAM GNCS project LESLIE.

## Footnotes

1 In the case of Continuous Time Markov Chains this construction is standard and the resulting DTMC is called *embedded chain.*

