## Supplementary Material for "A Conservative Approach for Describing Cancer Progression"

Nicolò Rossi, member, ETH Zürich
Nicola Gigante, member, Free University of Bozen-Bolzano
Nicola Vitacolonna, member, University of Udine
and Carla Piazza, member, University of Udine

June 6, 2022

### 1 Supplementary Figures

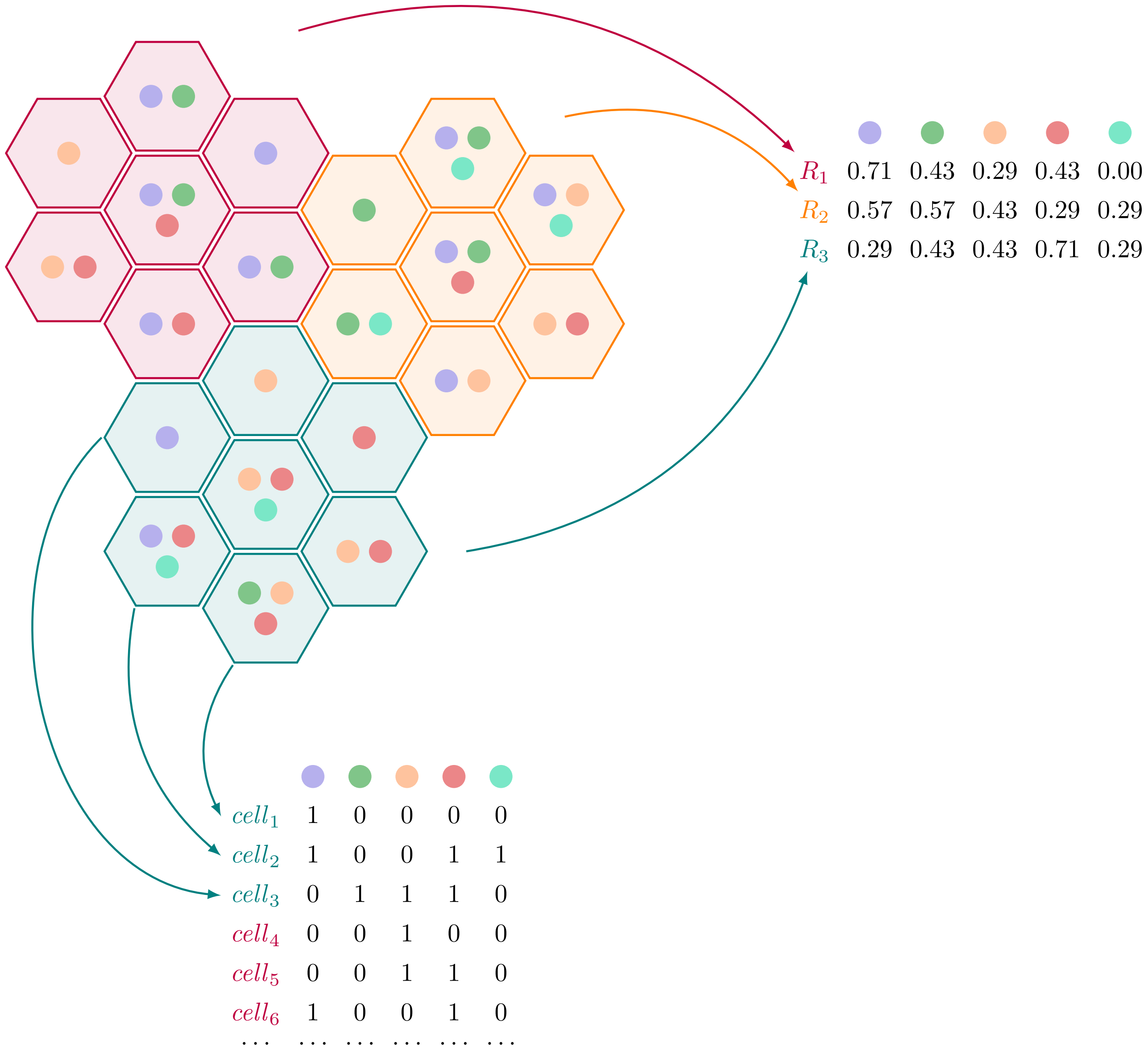


Figure 1: Bulk-regional and single-cell sampling.

### 1.1 Experiments on simulated data without siclonefit

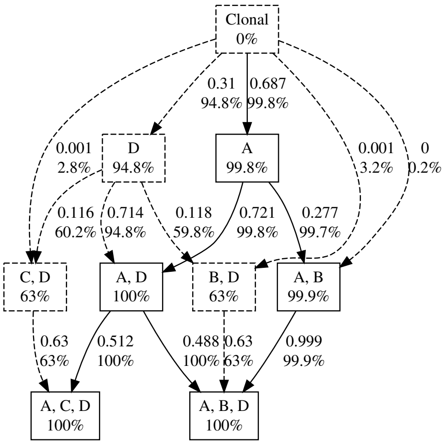


(a) $FP = 0.01$ and $FN = 0.05$.

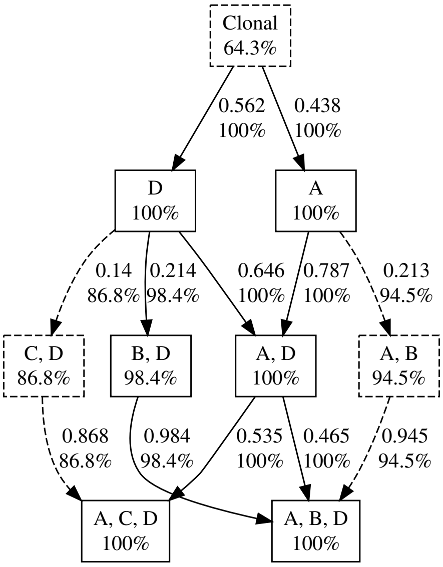


(b) $FP = 0.01$ and $FN = 0.15$.

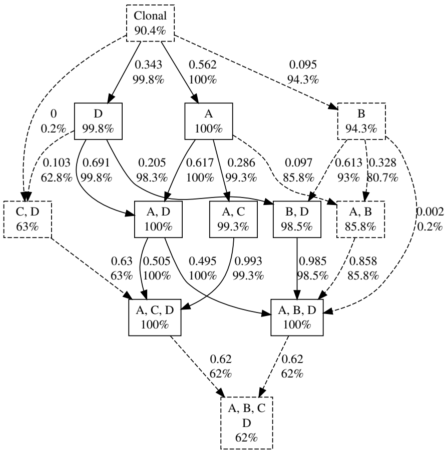


(c) $FP = 0.01$ and $FN = 0.15$.

Figure 2: Repetition 1. Examples without SiCloneFit preprocessing. Dashed components have a bootstrap probability less than 95%.

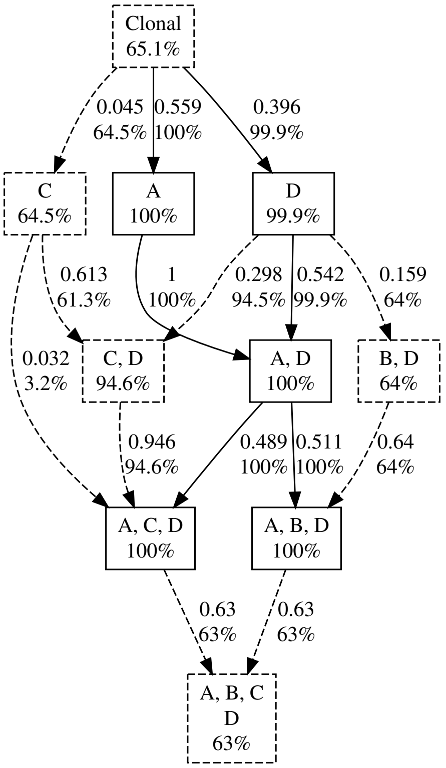


(a) $FP = 0.01$ and $FN = 0.05$.

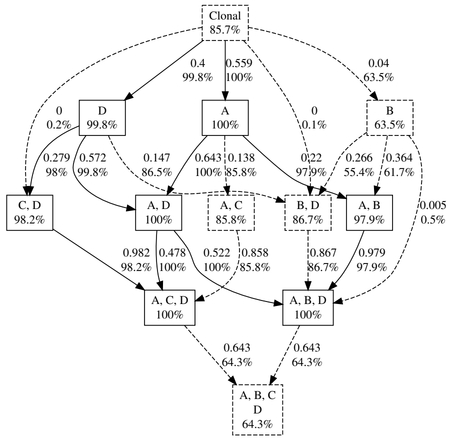


(b) $FP = 0.01$ and $FN = 0.15$.

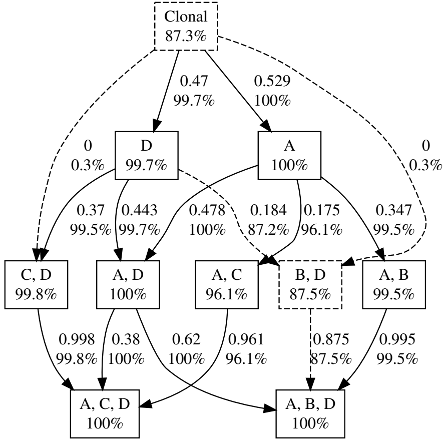


(c) $FP = 0.01$ and $FN = 0.15$.

Figure 3: Repetition 2. Examples without SiCloneFit preprocessing. Dashed components have a bootstrap probability less than 95%.

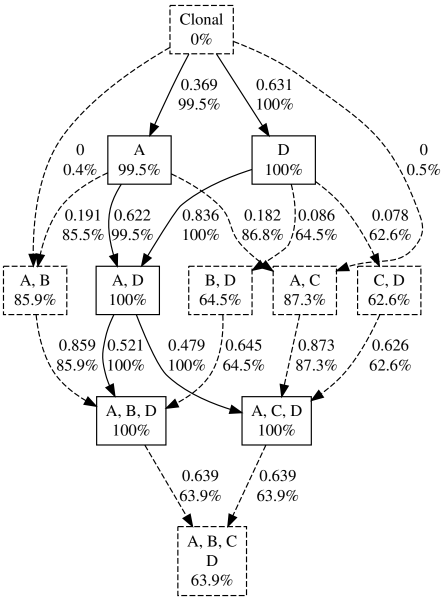


(a) $FP = 0.01$ and $FN = 0.05$.

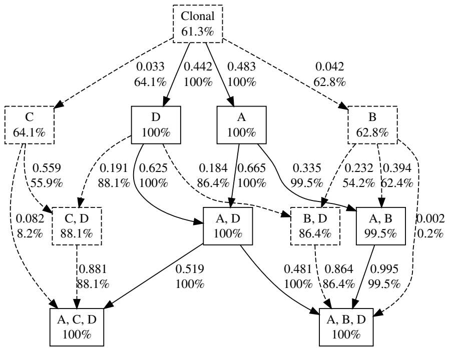


(b) $FP = 0.01$ and $FN = 0.15$.

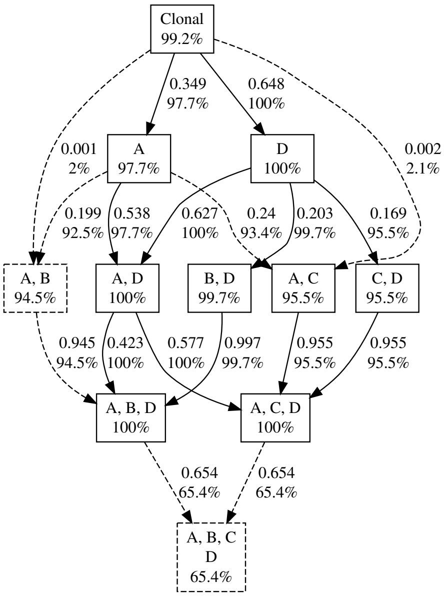


(c) $FP = 0.01$ and $FN = 0.15$.

Figure 4: Repetition 3. Examples without SiCloneFit preprocessing. Dashed components have a bootstrap probability less than 95%.

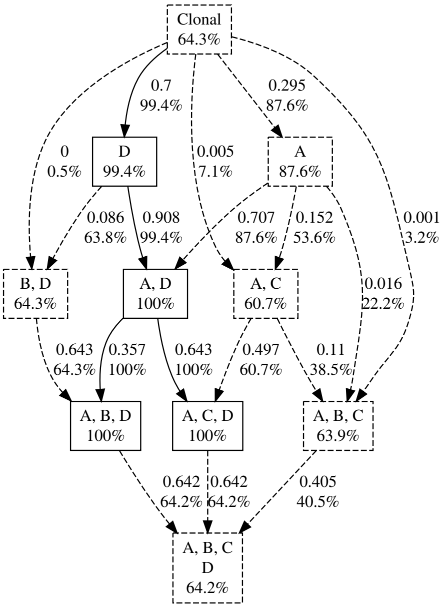


(a) $FP = 0.01$ and $FN = 0.05$.

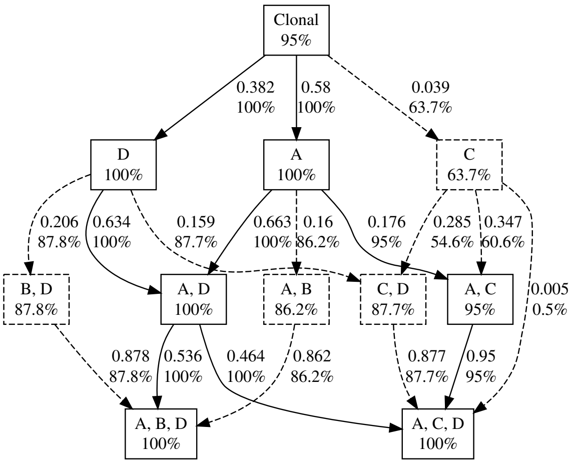


(b) $FP = 0.01$ and $FN = 0.15$.

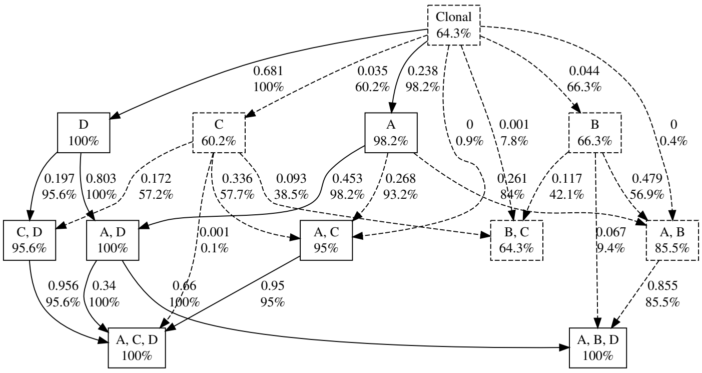


(c) $FP = 0.01$ and $FN = 0.15$.

Figure 5: Repetition 4. Examples without SiCloneFit preprocessing. Dashed components have a bootstrap probability less than 95%.

### 1.2 Experiments on simulated data with siclonefit

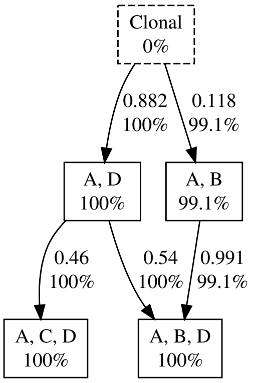


(a) $FP = 0.01$ and $FN = 0.05$.

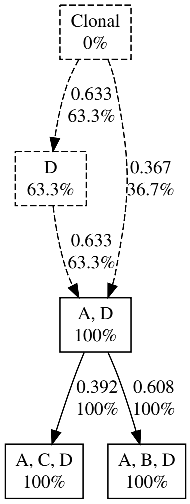


(b) $FP = 0.01$ and $FN = 0.15$.

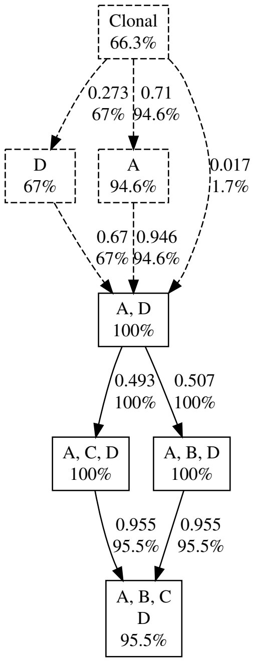


(c) $FP = 0.01$ and $FN = 0.15$.

Figure 6: Repetition 1. Examples without SiCloneFit preprocessing. Dashed components have a bootstrap probability less than 95%.

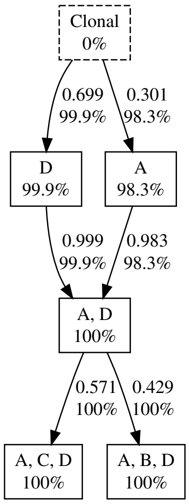


(a) $FP = 0.01$ and $FN = 0.05$.

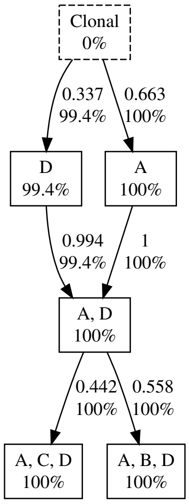


(b) $FP = 0.01$ and $FN = 0.15$.

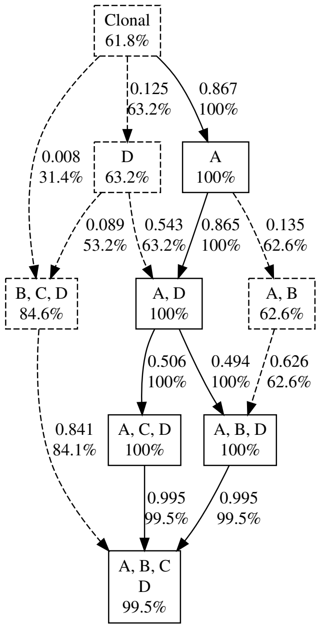


(c) $FP = 0.01$ and $FN = 0.15$.

Figure 7: Repetition 2. Examples without SiCloneFit preprocessing. Dashed components have a bootstrap probability less than 95%.

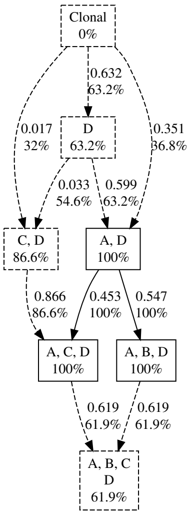


(a) $FP = 0.01$ and $FN = 0.05$.

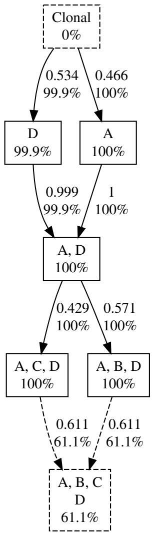


(b) $FP = 0.01$ and $FN = 0.15$.

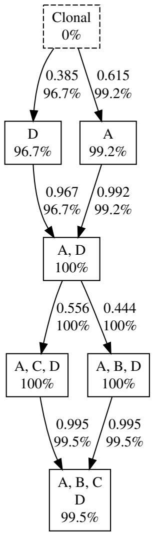


(c) $FP = 0.01$ and $FN = 0.15$.

Figure 8: Repetition 3. Examples without SiCloneFit preprocessing. Dashed components have a bootstrap probability less than 95%.

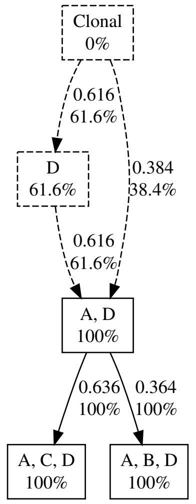


(a) $FP = 0.01$ and $FN = 0.05$.

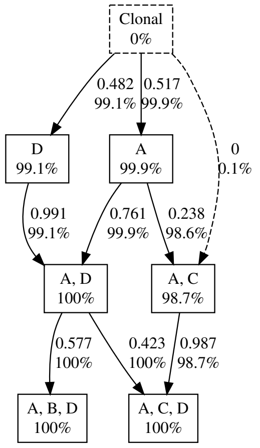


(b) $FP = 0.01$ and $FN = 0.15$.

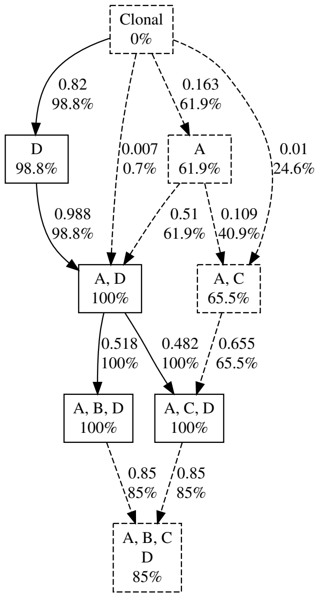


(c) $FP = 0.01$ and $FN = 0.15$.

Figure 9: Repetition 4. Examples without SiCloneFit preprocessing. Dashed components have a bootstrap probability less than 95%.

### 1.3 SCITE's results on the same datasets

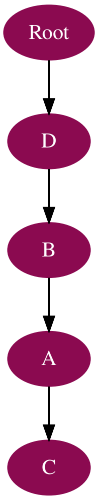


(a) $FP = 0.01$ and $FN = 0.05$.

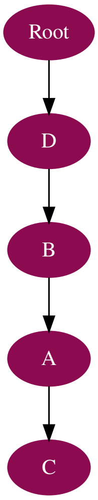


(b) $FP = 0.01$ and $FN = 0.15$.

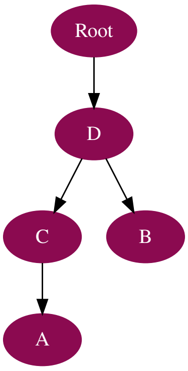


(c) $FP = 0.01$ and $FN = 0.15$.

Figure 10: Repetition 1. SCITE's results.

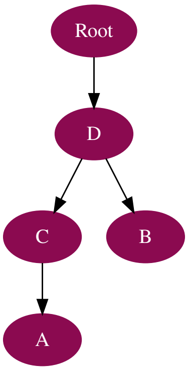


(a) $FP = 0.01$ and $FN = 0.05$.

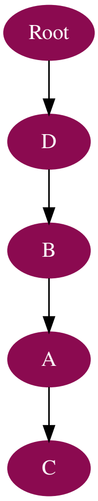


(b) $FP = 0.01$ and $FN = 0.15$.

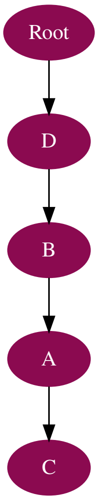


(c) $FP = 0.01$ and $FN = 0.15$.

Figure 11: Repetition 2. SCITE's results.

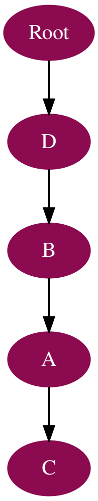


(a) $FP = 0.01$ and $FN = 0.05$.

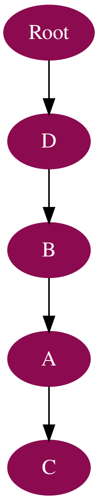


(b) $FP = 0.01$ and $FN = 0.15$.

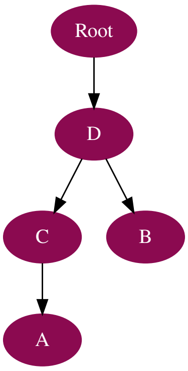


(c) $FP = 0.01$ and $FN = 0.15$.

Figure 12: Repetition 3. SCITE's results.

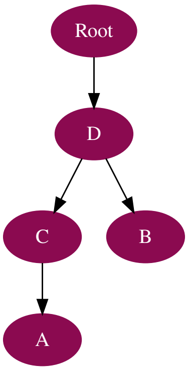


(a) $FP = 0.01$ and $FN = 0.05$.

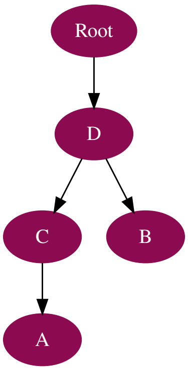


(b) $FP = 0.01$ and $FN = 0.15$.

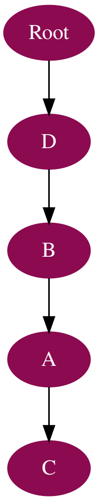


(c) $FP = 0.01$ and $FN = 0.15$.

Figure 13: Repetition 4. SCITE's results.
